# Insulin-Like Growth Factor 1 Receptor Regulates Breast Cancer Cell Adhesion through Beta-1 Integrin

**DOI:** 10.1101/2025.09.11.674989

**Authors:** Christopher A. Galifi, Elvan Dogan, Luis Fernandez Almansa, Krystopher Maingrette, Simran S. Shah, Joseph J. Bulatowicz, Karen Ebenezer, Amir K. Miri, Teresa L. Wood

## Abstract

**Introduction:** The insulin-like growth factor (IGF-1/IGF1R) pathway has been implicated in breast cancer aggressiveness; however, inhibition of this pathway has not been successful in clinical trials, indicating a lack of understanding about its role in TNBC metastasis. Recent studies have explored IGF1R involvement in integrin function and cancer cell adhesion dynamics. The goal of this study was to test the hypothesis that IGF1R itself regulates cancer cell adhesion.

**Methods:** We use MDA-MB-231 and Hs578T TNBC cell lines, siRNA-mediated knockdown, and adhesion assays to assess how IGF1R and integrin knockdowns impact cancer cell adhesion. Using xCELLigence E-plates, we quantify the effect of IGF-1 ligand stimulation versus IGF1R knockdown on functional cell adhesion. We also use HUVEC human endothelial cells to determine how IGF1R regulates adhesion to the endothelium.

**Results:** We found that IGF-1 stimulation increased MDA-MB-231 TNBC adhesion, which was reversed by the IGF1R tyrosine kinase inhibitor BMS-754807 and the ligand-dependent receptor internalization inhibitor dansylcadaverine. Unexpectedly, IGF1R knockdown also potently stimulated cell adhesion. Concomitant β1 integrin knockdown reversed the increased cell adhesion after both IGF-1 stimulation or IGF1R knockdown, indicating that the increased adhesion is β1 integrin dependent. This was also seen via immunocytochemistry when cells were seeded on fibronectin. Finally, inhibiting IGF1R signaling also reduced MDA-MB-231 cell adhesion to HUVEC endothelial cells.

**Discussion:** Both IGF-1 stimulation and IGF1R knockdown in TNBC cells promote cell adhesion, which seems paradoxical. However, the commonality of both interventions is removal of IGF1R from the cell surface, since IGF-1 stimulation causes IGF1R internalization and intracellular trafficking. Blocking IGF1R signaling using a tyrosine kinase IGF1R inhibitor preserves IGF1R on the cell surface. Thus, we propose a model whereby surface-bound IGF1R inhibits β1 integrin function and blocks cell adhesion. This model is supported further by our finding that treatment of MDA-MB-231 cells with dansylcadaverine, which inhibits ligand-mediated receptor internalization, blocked the effect of IGF-1 on adhesion. These findings may explain why selective IGF1R receptor antagonists, which downregulate IGF1R protein upon chronic administration, were unsuccessful in the clinical setting.

## Introduction

The role of the insulin-like growth factor type 1 receptor (IGF1R) in cancer is contentious, as preclinical studies of IGF1R function in cancer have shown both pro-tumorigenic and anti-tumorigenic functions (1, 2). While IGF1R inhibitors were developed and tested in clinical trials for the treatment of various cancers, no anti-IGF1R therapy has made it past phase 3. Moreover, no IGF1R inhibitor has made it past phase 2 for the treatment of breast cancer. For more information on the course of IGF1R inhibitors through clinical trials, please refer to our previous review on the topic (1). The poor results in clinical trials indicate that our knowledge of the contribution of IGF1R to tumor initiation and metastasis is incomplete; as such, there is considerable interest in investigating the molecular mechanisms of crosstalk between IGF1R and other receptor or signaling proteins.

IGF1R is a heterotetrametric tyrosine kinase receptor with high homology to the insulin receptor (INSR) (3). A recent phosphoproteomic analysis of IGF1R versus INSR function in brown preadipocytes found that IGF1R signaling preferentially promoted cell proliferation, whereas INSR signaling was pro-metabolic (4). These results are consistent with numerous studies showing a prominent role for the IGF1R in proliferation and survival of cancer cells(5–7). In addition, several studies revealed that IGF1R has a prominent function in cell adhesion, (for recent review, see (1)).

In prior studies, we generated mouse lines of Wnt1-driven mammary tumors, a model of basal-like TNBC, with decreased IGF1R function, either via mammary epithelial-specific gene deletion or expression of a dominant-negative IGF1R transgene (8–10). Loss or inhibition of IGF1R signaling resulted in decreased tumor latency and increased incidence of lung metastases (8, 9). We also observed that inhibiting IGF1R function in these mouse models caused dysregulation of tumor cell adhesion (9). Conversely, studies from the O’Connor laboratory revealed that cancer cell adhesion promotes signaling through the IGF1R and permits an aggressive phenotype (11). Thus, in addition to its established functions in cell proliferation and survival (12, 13). these findings support functions for the IGF1R in regulating cell adhesion, which when dysregulated, contributes to anchorage-independent growth, cell invasion and metastasis.

Precisely how IGF signaling and the IGF1R regulate cell adhesion is unclear; however, emerging research suggests IGF1R crosstalk with integrins, the main adhesion receptors responsible for adherence to the extracellular matrix (ECM) (14, 15). Colocalization between IGF1R and integrins have been reported (14, 16) including the β1 subunit (17, 18). Recently, β1 integrin was found to cooperate with IGF1R to promote its function and relocalization in the cell (18). Investigators in an early study found that IGF-1 induced an association between IGF1R and receptor for activated C kinase 1, RACK1, (19), a scaffolding protein capable of binding to the cytoplasmic domain of β1 integrin (20). These previous studies focused on the role of integrins in modulating IGF1R function in cancer (14, 18, 21); however, we hypothesized that the interaction is reciprocal such that IGF1R regulates cancer cell adhesion during metastatic progression through its interaction with integrins. Although integrin signaling is a well-known mediator of cell to ECM adhesion, the role of IGF1 and the IGF1R on integrin function in tumor cell – ECM adhesion is not well understood. Our goal in this study was to investigate how IGF-1 and IGF1R regulate β1 integrin.

## Materials and Methods

### Clinical Data Mining and Analysis

The data generated from 2509 patients within the METABRIC project (22, 23) was used in this investigation. These data were accessed through Synapse (https://synapse.sagebase.org), including normalized expression data and clinical feature measurements. The associated expression Z scores were downloaded from cBioPortal (24–26) (https://www.cbioportal.org/). IGF1R expression was queried, restricting genomic profiles to mRNA expression z-scores relative to all samples with a z-score threshold of 1. Patients above this threshold were classified as high IGF1R and those below were classified as low IGF1R. Custom groups were developed from high and low IGF1R patient data. Comparative analyses between high and low IGF1R groups were performed using the portal’s built-in group comparison tool. Fibronectin and β1 Integrin expression was mined for within these high and low IGF1R groups.

### Cell Culture and Transfection

Cell culture procedures are identical to those from our recent manuscript (27). Cell lines were purchased from ATCC (Manassas, VA) and were authenticated in the past year by ATCC. The cell lines are tested for mycoplasma contamination twice annually. MDA-MB-231(28) (RRID:CVCL_0062) and Hs578T(29) (RRID:CVCL_0332) cells were maintained in DMEM (Thermo Fisher, Waltham, MA) with 10% fetal bovine serum (FBS) and 1% penicillin/streptomycin (P/S) at 37°C and 5% CO_2_. Hs578T media was supplemented with 0.01 mg/ml human insulin per ATCC’s instructions during maintenance culture; insulin-supplemented media was replaced with insulin-free media during experiments. Media changes or passages were performed 2-3 times per week. Cells were seeded at a density of 5x10^5^ per well in 6 well plates one day prior to transfection. In all cases, siRNA was transfected at a final concentration of approximately 8 nM in each well with 7.5 µl Lipofectamine RNAiMAX reagent per well (Thermo Fisher #13778030) per the manufacturer’s instructions. Cells were collected on day 3 post-transfection for xCELLigence. Plasmids for overexpression studies were transfected via Lipofectamine 3000 reagent (Thermo Fisher # L3000008), using 7.5µg of plasmid per well, and cells were collected for xCELLigence 2 days post-transfection. All siRNAs, plasmids, and qPCR primers used are listed in Table 1. We used these drugs for inhibition studies at the following concentrations: IGF1R/INSR inhibitor BMS-754807 (SelleckChem, Houston, TX), 10 nM; MEK inhibitor U0126 (Cell Signaling, Danvers, MA), 10 µM; PI3K inhibitor LY294002 (Cell Signaling), 50 µM; inhibitor of ligand-dependent receptor internalization dansylcadaverine (MedChemExpress, Monmouth Junction, NJ), 100 µM as previously described (30).

**Table 1.**
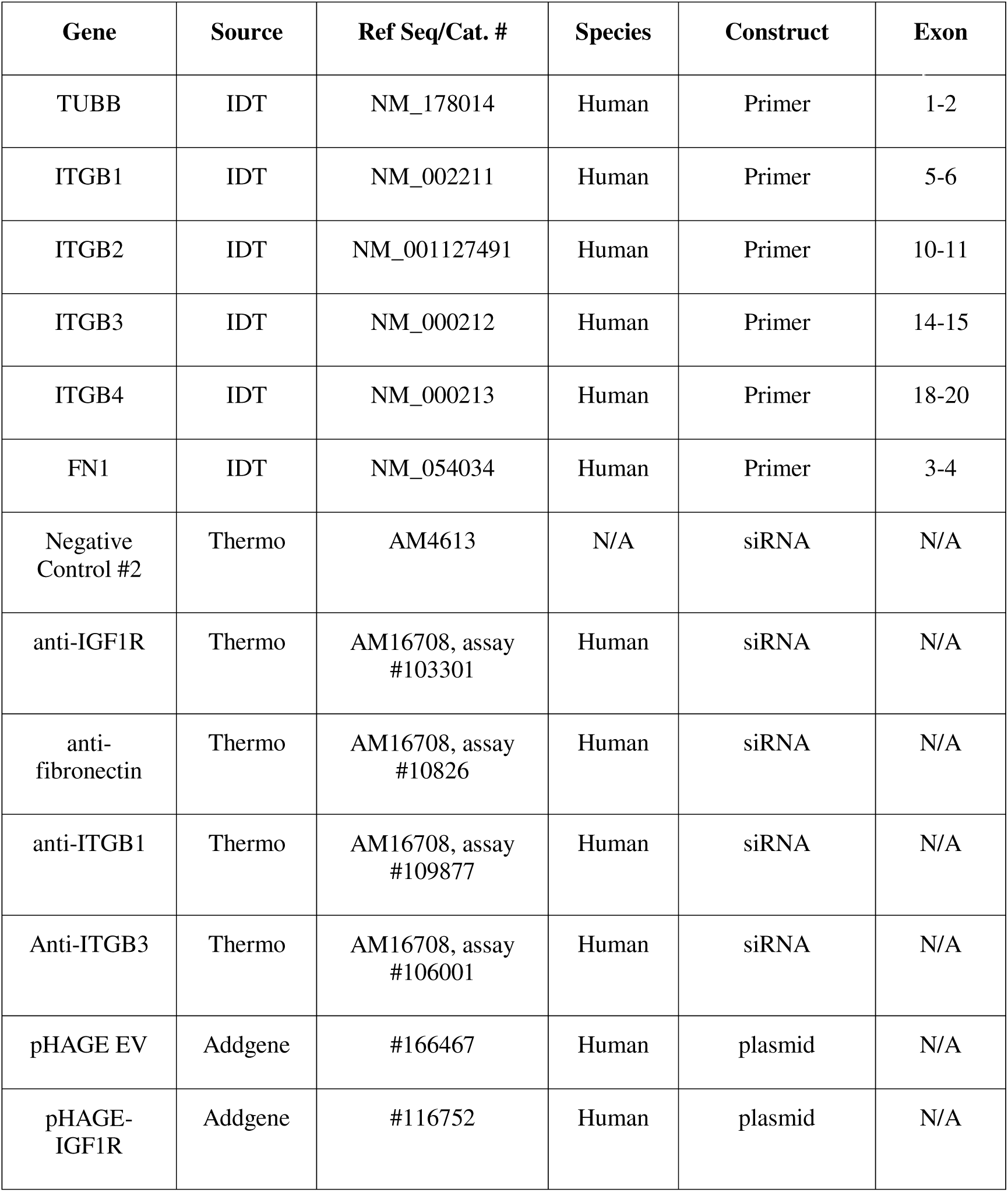
List of all primers, siRNA, and plasmids used in this study.

### xCELLigence

16-well xCELLigence E-plates were purchased from Agilent (cat. #300600890). Just prior to collection, cells were serum-starved in DMEM base media with or without drug treatments or vehicle controls as indicated in the figure legends. Cells were detached in accutase (Sigma-Aldrich #A6964) and collected in equal parts DMEM supplemented with the indicated treatments. Mastermixes of 3x10^5^ cells/ml were made per treatment group using treatment-supplemented DMEM, and IGF-1 was supplemented while cells were in suspension just prior to seeding xCELLigence E-plates. 3x10^4^ cells were seeded per well. Cell index measurements were made every 3 minutes for 3 hours during adhesion experiments.

### RT-qPCR

To assess basal levels of integrin expression in each cell line, 5x10^5^ cells were seeded in 6 well format in DMEM, 10% FBS, and 1% P/S and incubated at 37°C overnight. On the next day, cells were scraped and collected for RNA isolation. For validation experiments, cells were collected either through this manner or from residual cells left over from xCELLigence experiments. RNA was extracted using the Qiagen RNeasy Mini Kit (cat. #74104). cDNA was synthesized via the BioRad iScript cDNA synthesis kit (cat. #170-8891) with 500 ng of RNA. cDNA was diluted 1:5 in sterile water, and qPCR was performed using the iTaq Universal SYBR Green Supermix (BioRad #1725124). Relative expression values were calculated using the qgene macro (31) and multiplied by 10^3^ to increase numbers above decimal fractions for easier interpretation. Primers used are listed in Table 1.

### Protein Isolation, SDS-PAGE and Western Blot

Our general western blot workflow is identical to our recent report (27). Protein samples in Laemmli buffer were electrophoresed through NuPAGE 4-12% polyacrylamide gels (Thermo Fisher), then transferred to nitrocellulose membranes (Amersham, UK). Membranes were blocked with 5% milk in tris buffered saline with 0.1% Tween 20 (TBS-T) for 1 hour at room temperature prior to incubation with each primary antibody. Membranes were incubated with primary antibodies in 5% bovine serum albumin (BSA) in TBS-T for either 1 hour at room temperature or overnight at 4°C. Antibodies used include anti-IGF1Rβ (Cell Signaling #3027, RRID:AB_2122378), anti-phospho-IGF1R/IR (Cell Signaling #3024, RRID:AB_331253), anti-β-tubulin (Cell Signaling #2146, RRID:AB_2210545), anti-AKT (Cell Signaling #9272, RRID:AB_329827), anti-phospho-AKT Ser473 (Cell Signaling #4060, RRID:AB_2315049), anti-phospho-ERK1/2 (Cell Signaling #4370, RRID:AB_2315112), anti-ERK1/2 (Cell Signaling #9102, RRID:AB_330744), anti-phospho-FAK (Tyr397) (Thermo #44-625G, RRID:AB_2533702), anti-FAK (Cell Signaling #3285, RRID:AB_2269034), anti-fibronectin (Cell Signaling #26836, RRID:AB_2924220), anti-beta-1 integrin (Cell Signaling #34971, RRID:AB_2799067), anti-phospho-Talin (Cell Signaling #5426, RRID:AB_10695406), anti-Talin-1 (Cell Signaling #4021, RRID:AB_2204018, anti-cofilin (Cell Signaling #5175, RRID:AB_10622000), anti-PP2A (Cell Signaling #2038, RRID:AB_2169495), anti-RACK1 (Cell Signaling #5432, RRID:AB_10705522), and anti-GAPDH (Cell Signaling #5174, RRID:AB_10622025). Probed membranes were then incubated with horseradish peroxidase-conjugated secondary antibodies (Jackson ImmunoResearch, West Grove, PA), diluted 1:5000 in 5% milk in TBS-T, for 1 hour at room temperature prior to analysis via enhanced chemiluminescence (Revvity). Western blot quantification was performed using ImageJ (32). The selection tool was used to outline sample bands within a fixed area per protein target, and the measurement tool was used to determine average pixel intensity. Background signal was removed using the subtract tool in ImageJ. All raw blots and ROIs used will be uploaded to our OSF repository.

### Cell Adhesion and Immunostaining

To validate the cell adhesion experiment conducted with the xCELLigence system, a fluorescence assay was performed through an endothelial barrier using Lab-Tek™ II chamber slides (Thermo Fisher #154534). Glass surfaces were pre-coated with either poly-D-lysine (PDL; Millipore P1024-100MG)) or 50 µg/mL fibronectin (Sigma-Aldrich # 11051407001) for 1 h at RT and washed in PBS. 1x10^4^ cells were grown on pre-coated slides for 3 hours in base DMEM with or without 10 nM BMS. After 3 hours, MDA-MB-231 cells were stimulated with 10 nM IGF-1 for 1 hour prior to fixation with 4% paraformaldehyde in PBS for 15 minutes at room temperature. Cells were permeabilized with 0.1% Triton X-100 in PBS for 30 minutes at room temperature, then blocked in 1% BSA in PBS for 1 hour at room temperature. Cells were then incubated with primary antibodies in 1% BSA in PBS overnight at 4°C. We used Integrin β1 (D6S1W) Rabbit mAb (CST #34971) and Goat anti-Rabbit IgG (H+L) Cross-Adsorbed Secondary Antibody, Alexa Fluor™ 488 (Invitrogen #1851447). Cells were then incubated with secondary antibody diluted 1:500 in PBS for 1 hour at room temperature prior to coverslip mounting with ProLong Gold mountant (Invitrogen #2936373). After imaging, the data were quantified in two ways: (i) the average cell attachment area was measured to determine how each treatment influenced the size of individual attachment sites, and (ii) the total number of attached cells was counted under 4× magnification to evaluate overall cell attachment to the substrates.

### Adherence to HUVEC Endothelial Cells

A fluorescence assay was performed through an endothelial barrier using Lab-Tek™ II chamber slides (Thermo Fisher #154534). Glass surfaces were pre-coated with 50 µg/mL fibronectin (Millipore Sigma #1108093800) for 1 h at RT and washed in PBS. Human umbilical vein endothelial cells (HUVECs; ATCC #CRL-1730 ™) were seeded at 1×10 cells/well in EGM-2 medium and permitted to culture for 24 h to establish a confluent monolayer. MDA-MB-231 cells were suspended-stained using BioTracker™ 490 Green Cytoplasmic Membrane Dye (Sigma-Aldrich # SCT106) at the manufacturer-recommended concentration and cotreated with 100 nm IGF1R/INSR inhibitor BMS-754807 (SelleckChem, Houston, TX) for 30 min in an incubator (37°C, 5% CO □). Cells were then centrifuged and washed three times with PBS to remove any residual dye and BMS. The culture medium was replaced with a 1:1 mixture of DMEM (for the growth of MDA-MB-231) and EGM-2 (for the growth of HUVEC) just prior to seeding stained MDA-MB-231 cells onto the HUVEC monolayer in order to culture both cell types using the intravasation assay. 2.5×10□ of stained tumor cells were carefully added to each well. After 24 h of incubation and PBS washing step, slides were imaged using both brightfield and green fluorescence channels to perform fluorescence intensity analysis using ImageJ.

### Statistics

Biological replicates of cell lines were established either from separate thaws of cell lines run in an experiment on the same day, or from separate passages of cell lines run in experiments on separate days. Statistical analysis was performed using Prism Graphpad. One-way analysis of variance (ANOVA) with post-hoc Tukey’s or Šidák multiple comparison tests to determine statistical significance between multiple groups or Dunnett’s test for statistical significance between control and >1 experimental groups with α set to 0.05 as the threshold for significance. Two-tailed t-tests were used to assess significance in experiments with two groups. For the validation of IGF1R overexpression, one-tailed t-test was used, as an increase in IGF1R protein was expected.

## Results

Since our previous findings revealed that low IGF1R expression in human breast tumors correlates with worse clinical prognosis and reduced IGF1R also increased metastasis in a mouse model of TNBC and altered tumor cell adhesion dynamics (9, 10), we investigated whether integrin expression is altered in breast cancers with changes in expression of IGF1R. Prior investigators found strong associations between IGF1R and β1 integrin (17, 18, 33), so we began by searching for β1 integrin expression, as well as its ligand fibronectin, in human breast cancers by mining the cancers by mining the METABRIC dataset (22, 23) using cBioPortal (24–26). In breast tumors with low expression of IGF1R, we found significantly elevated expression of β1 integrin, and fibronectin supporting a link between IGF1R expression and β1 integrin function in breast cancers (Figure 1A-B).

**Figure 1.**
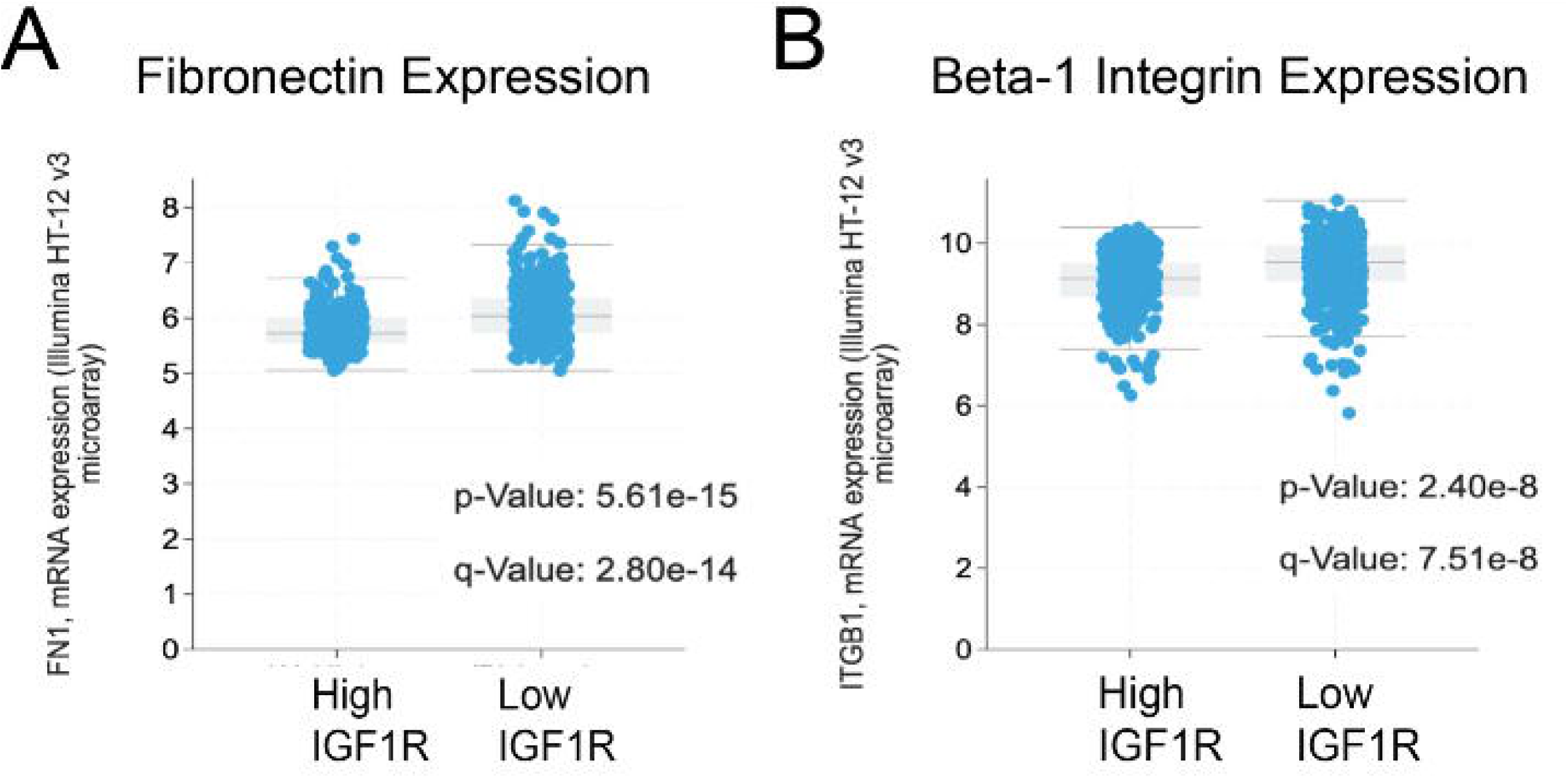
Breast cancer patients with low IGF1R expression have increased expression of extracellular matrix-cell adhesion molecules. METABRIC data analysis for A) Fibronectin and B) β1 integrin in patient tumors with High IGF1R (IGF1R z-score > 1) and Low IGF1R (IGF1R z-score < 1). All extracellular matrix-cell adhesion molecules show an inverse relationship with IGF1R expression. Statistical Analysis preformed using Student t-test.

We were then interested to further investigate a mechanistic relationship between IGF1R, cell adhesion and integrin function in breast cancer. To test how IGF-1 alters adhesion of TNBC cells, we used MDA-MB-231 and Hs578T human TNBC cell lines that both carry Ras mutations. We analyzed changes in cell adhesion using an xCELLigence real-time assay in the presence or absence of stimulation with 10 nM IGF-1. IGF-1 increased cell adhesion by twofold in MDA-MB-231 cells (Figure 2A, C) but did not significantly affect baseline Hs578T adherence (Figure 2B, D). Treatment with an IGF1R/INSR inhibitor (BMS-754807) abrogated the effect of IGF-1 on cell adhesion in the MDA-MB-231 cells (Figure 2A, C). Since xCELLigence E-plates are uncoated so devoid of a substrate that activates integrins, these data suggest that IGF-1 can promote adhesion of MDA-MB-231 cells without an additional ECM stimulus.

**Figure 2.**
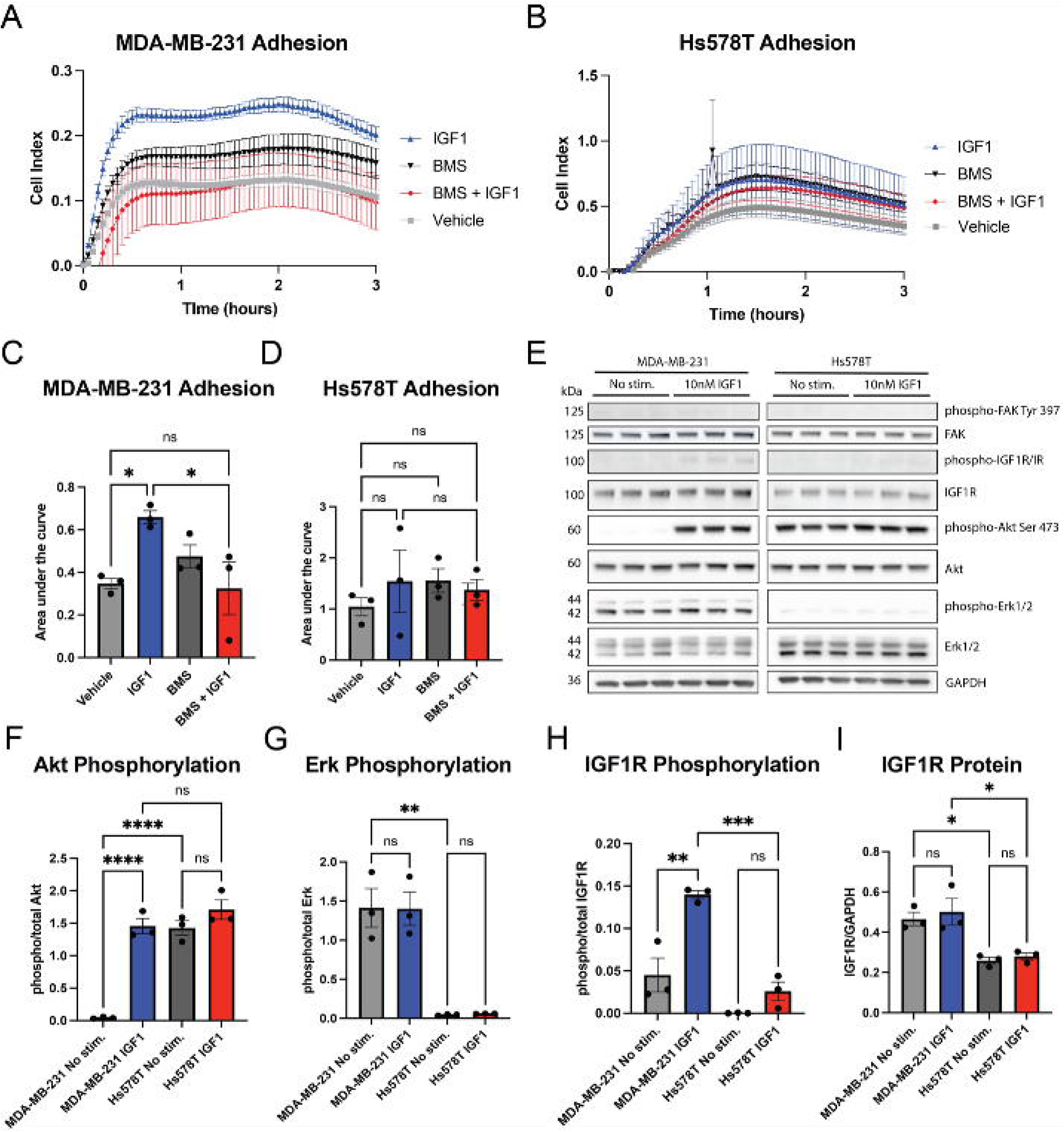
IGF-1 increases adherence and AKT phosphorylation in MDA-MB-231 cells. A,B) Representative tracings showing MDA-MB-231 (A) and Hs578T (B) cell adhesion over time using xCELLigence assays. C,D) Area under the curve for n=3 separate experiments for each cell line was analyzed for significance via one-way ANOVA and Šidák multiple comparison post hoc test. E) Western blot showing effect of IGF-1 stimulation on downstream signaling in MDA-MB-231 and Hs578T cells. F-I) Graphs showing quantification of data in (E) analyzed by one-way ANOVA and Tukey’s post hoc test. *p<0.05, **p<0.005, ***p<0.0005, ****p<0.0001.

IGF-1 signaling through IGF1R can activate both the ERK and AKT pathways (34). However, the role of each pathway in regulating cell adhesion is poorly understood. To identify pathway activation associated with IGF-1-induced adhesion in MDA-MB-231 cells, we assessed AKT and ERK1/2 phosphorylation as well as that of IGF1R. We also measured phosphorylation of focal adhesion kinase (FAK), since this protein is critical for the organization of integrins (35) and is implicated in cancer progression (36). Interestingly, IGF-1 potently stimulated the AKT pathway in MDA-MB-231 cells but not in Hs578T cells due to constitutive activation of AKT in this cell line (Fig. 2E, F. In MDA-MB-231 cells, ERK1/2 were both phosphorylated at baseline, and IGF-1 stimulation had no additional effect on their phosphorylation (Figure 2E, G). In Hs578T cells, levels of P-ERK1/2 were low regardless of IGF-1 treatment (Fig. 1E, G). Surprisingly, FAK phosphorylation was low to undetectable with or without IGF-1 stimulation in both cell lines (Fig. 2E) suggesting that IGF-1 may promote adhesion in a FAK-independent manner in the MDA-MB-231 cells.

IGF-1 increased phosphorylation of IGF1R in MDA-MB-231 cells (Figure 2E, H), but consistent with the low adhesion response to IGF-1, IGF-1 stimulation failed to significantly increase phosphorylation of IGF1R in Hs578T cells (Figure 2E, H). Since levels of IGF1R were two-fold lower in Hs578T cells compared to MDA-MB-231 cells (Figure 2E, I), we overexpressed IGF1R in this cell line to determine if the insensitivity to IGF-1 was due to low IGF1R levels. Transfection of an IGF1R overexpression plasmid significantly increased levels of IGF1R protein in the Hs578T cells (Supplemental Figure 1A-B). However, IGF-1 failed to increase adhesion despite heightened expression of IGF1R in the transfected Hs578T cells (Supplemental Figure 1C-D), suggesting resistance to IGF-1 in this TNBC line is due to factors other than receptor expression. Based on these findings, we proceeded with the MDA-MB-231 cell line to further understand the interaction of IGF1R with integrins in cell adhesion.

Since IGF-1 stimulation induced a sustained increase in adhesion with a plateau at approximately 30 minutes in the MDA-MB-231 cells (Figure 2A), we analyzed phosphorylation of IGF1R and downstream targets over a similar timeframe. Both IGF1R and AKT phosphorylation increased from 5-60 min with IGF-1 stimulation, consistent with IGF-1-induced adhesion (Figure 3A-C). Similar to our previous findings, ERK1/2 phosphorylation was unchanged in the MDA-MB-231 cells over time (Figure 3A, D), consistent with constitutive activation of the pathway due to a known KRAS mutation in these cells (37). Interestingly, activation of Talin, a marker of integrin activation, was also unchanged following IGF-1 stimulation through 60 min, suggesting that it is not involved in IGF-1 mediated cell adhesion (Figure 3A, E).

**Figure 3.**
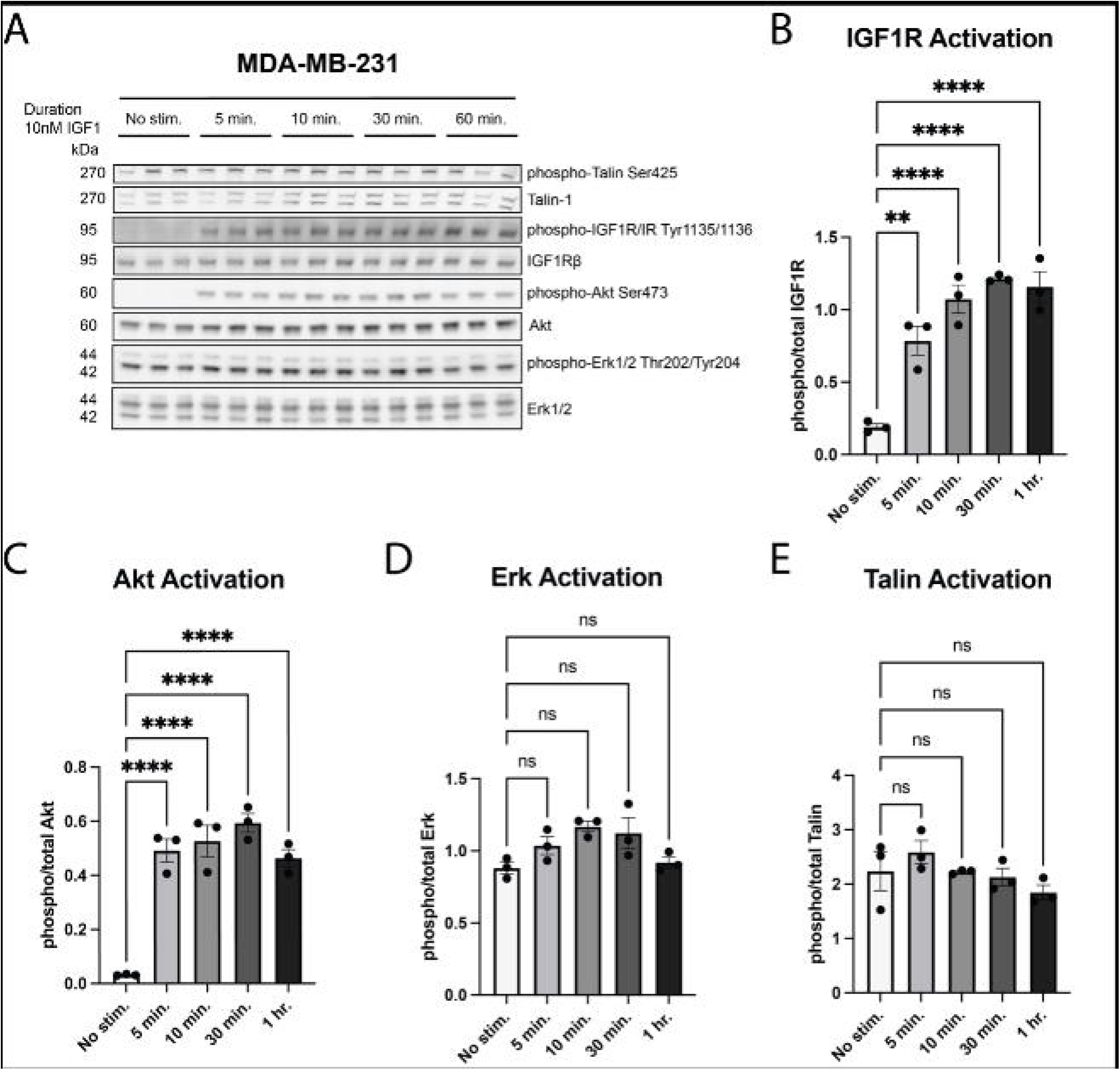
IGF-1 activates IGF1R and AKT in MDA-MB-231 cells, but Talin is constitutively active and remains unaffected by IGF-1 stimulation. A) MDA-MB-231 cells were treated with IGF-1 at the indicated timepoints and downstream signaling targets were analyzed via western blot. B-D) Phospho-total levels were plotted to determine activation of IGF1R (A), AKT (B), ERK1/2 (C), and TALIN (D) across timepoints, assessed via one-way ANOVA and Dunnett’s post hoc test. **p<0.005, ****p<0.0001.

Previously, we reported that IGF1R internalization was necessary for prolonged AKT stimulation, specifically in glial progenitor cells (30). In this prior study, dansylcadaverine (DC), an inhibitor of transglutaminase-mediated receptor internalization, prevented both receptor internalization and AKT phosphorylation in a rat glial progenitor cell line. DC similarly inhibited IGF1R internalization in Chinese hamster ovary cells in another study (38); however, in these cells, DC did not prevent activation of AKT or of insulin receptor substrate-1 (IRS-1), the primary signal transducer of the AKT pathway through IGF1R. This suggests that the impact of IGF1R internalization on intracellular signaling is cell-type dependent. To determine the effect of inhibiting IGF1R internalization on MDA-MB-231 adhesion over time, cells were pretreated with DC before stimulating with IGF-1. Strikingly, inhibition of IGF1R internalization completely abrogated IGF-1-dependent adhesion (Figure 4A, C). IGF-1 in the presence of DC had no effect on either IGF1R phosphorylation (Figure 4B, D) or AKT phosphorylation in MDA-MB-231 cells (Figure 4B, E). ERK1/2 activation was unchanged after IGF-1 stimulation regardless of DC administration (Figure 4B, F). These results suggest that IGF1R internalization is sufficient to promote IGF-1-dependent cell adhesion in MDA-MB-231 cells independently of its phosphorylation or activation of AKT.

**Figure 4.**
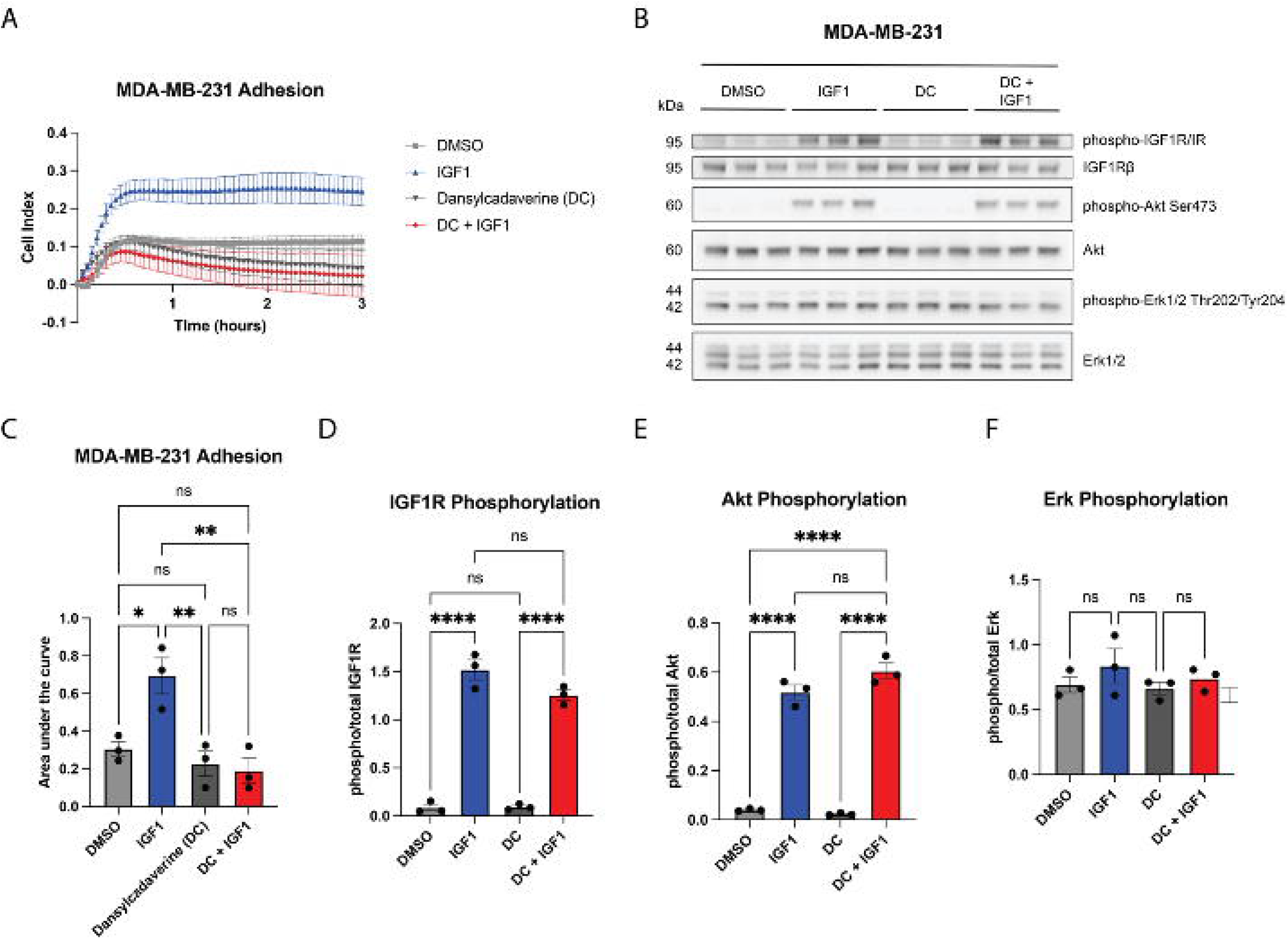
IGF-1 promotes adhesion in MDA-MB-231 TNBC cells that is dependent on IGF1R internalization. A) Representative tracings from xCELLigence real-time assay showing adhesion of MDA-MB-231 cells treated with IGF-1 in the presence or absence of dansylcadaverine (DC), an inhibitor of receptor internalization. Quantification of n=3 independent experiments is shown in (C) Cell adhesion was quantified via area under the curve, one-way ANOVA and Tukey’s post-hoc test. B) The effects of DC on IGF-1-dependent signaling were assessed via western blot. D-F) Graphs showing quantification of (B) for phospho-total levels of of IGF1R (D) or AKT (E) and ERK (F) assessed via one-way ANOVA and Tukey’s posthoc test. *p<0.05, **p<0.005, ****p<0.0001.

Since we established that IGF1R internalization is a main contributing factor to IGF-1-dependent adhesion, we were then interested in determining the role of integrins in this process. We first assessed mRNA expression of integrin subunits β1-4 to determine which were most likely to be involved in adhesion in the two TNBC cell lines (Supplemental Figure 2A-B). In both MDA-MB-231 and Hs578T cells, β1 integrin was the most highly expressed of the beta subunits. This is consistent with the literature, as most integrin heterodimers are composed of one β1 subunit paired with an α subunit (39). Both cell lines robustly expressed β1 integrin protein (Supplemental Figure 2C-D). High expression of β1 integrin in cancer is unsurprising, as expression of this subunit along with β3 integrin is a marker of the epithelial-mesenchymal transition, a hallmark of aggressive cancers (40). In addition, β1 integrin often heterodimerizes with α5 integrin to form one of the primary fibronectin receptors (41). Since fibronectin expression is strongly correlated with IGF1R expression in human breast tumors (Figure 1), we analyzed fibronectin protein expression in the TNBC cell lines (Supplemental Figure 2C, E). Interestingly, fibronectin expression was only detectable in the Hs578T cell line. Finally, since crosstalk between β1 integrin and IGF1R was found to be mediated by PP2A and RACK1 in a previous study (42), we assessed expression levels of these proteins in the two cell lines (Supplemental Figure 2C, F, G). Both cell lines expressed similar levels of PP2A and RACK1.

Because β1 integrin was the predominant integrin subunit expressed by MDA-MB-231 cells, we performed siRNA knockdown of ITGB1 mRNA to determine if this integrin is necessary for IGF1-dependent adhesion in the MDA-MB-231 tumor cells. First, we confirmed the efficacy of siRNA-mediated knockdown by analyzing expression of β1 integrin protein (Supplemental Figure 3A, B). We then analyzed adhesion of the cells after knockdown of β1 integrin. IGF-1 stimulated cell adhesion in control transfected cells but failed to enhance adherence after knockdown of β1 integrin (Figure 5A-B) indicating that β1 integrin is necessary for IGF-1-dependent adhesion in MDA-MB-231 cells. We also attempted to knock down β3 integrin in these cells but were unsuccessful possibly due to low baseline expression of this subunit (Supplemental Figure 3A, C).

**Figure 5.**
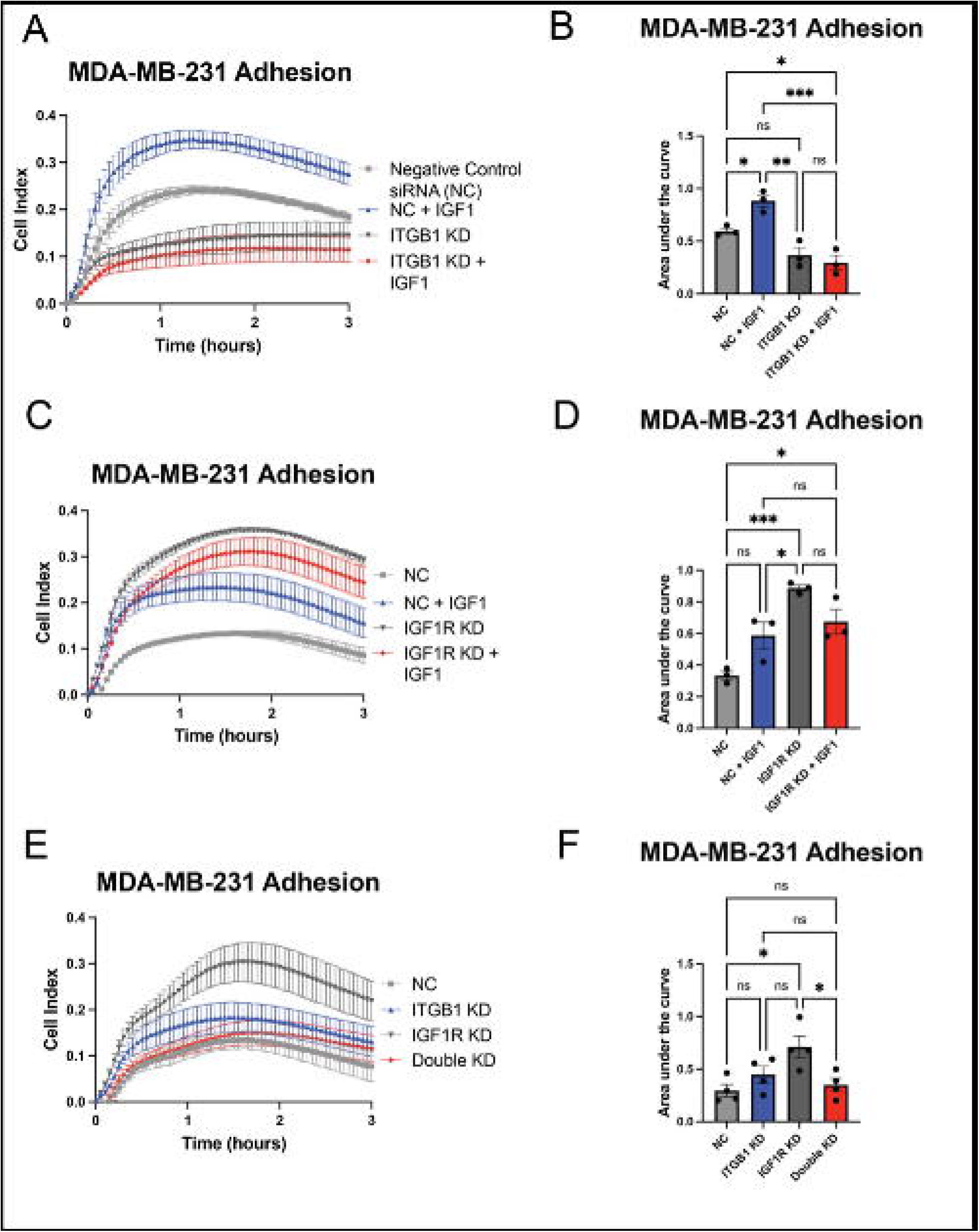
IGF-1-mediated increases in adhesion are dependent on β1 integrin. A,C,E) Representative tracings from xCELLigence adhesion assays after siRNA-mediated β1 integrin knockdown (A), IGF1R knockdown (C) or both (E) in MDA-MB-231 cells with or without treatment with IGF-1. B,D,F) Quantifications of n=3 independent experiments for each knockdown assessed via one-way ANOVA and Tukey’s posthoc test. *p<0.05, **p<0.005, ***p<0.0005

Based on these data, we hypothesized a mechanism whereby IGF-1 binding to IGF1R induces receptor internalization promoting β1 integrin function at the cell surface. As such, we predicted that IGF1R knockdown without IGF-1 binding would also promote cell adhesion by removing IGF1R from the cell surface. Consistent with this, we found that siRNA-mediated IGF1R knockdown enhanced adhesion in the MDA-MB-231 cell line (Supplemental Figure 3A, C; Figure 5C, D). The increase in cell adhesion was statistically greater than both baseline adhesion in serum-starvation conditions as well as in IGF-1-stimulated conditions. Consistent with our hypothesis, the increase in adhesion after IGF1R knockdown was also β1 integrin-dependent, as co-transfection of anti-IGF1R and anti-β1 integrin siRNAs reduced MDA-MB-231 cell adhesion back to baseline (Figure 5E, F). These results indicate that, paradoxically, both IGF1 stimulation and IGF1R knockdown promote TNBC adhesion; however, we propose that the mechanism in both cases is due to removal of IGF1R from the cell surface.

Thus far in the real-time adhesion assays, we used xCELLigence E-plates that were uncoated and lacked substrate capable of activating integrins; as such, the data reflect the ability of IGF-1 or anti-IGF1R siRNA to increase integrin function in the absence of substrate. Integrin-activating substrates such as fibronectin can provide a physical platform for cancer cells to attach and migrate (43). To determine if IGF-1-dependent adhesion was altered in the presence of integrin binding, we repeated siRNA-mediated knockdown of both IGF1R and β1 integrin using plates coated with 0.05 mg/ml fibronectin, a major ECM component and ligand for α5β1 integrin. Fibronectin substrate abolished the increased adherence of MDA-MB-231 cells treated with IGF-1 such that their adherence was similar to serum-starved control cells (Figure 6A, B). Fibronectin induced a sharp adherence peak at approximately 1 hour, which then diminished over time in both the β1 integrin and IGF1R knockdown experiments (Figure 6A, C). β1 integrin knockdown significantly flattened this peak in both the IGF-1-stimulated and unstimulated groups (Figure 6A, B), suggesting that β1 integrin is critical for binding to fibronectin in MDA-MB-231 cells. This was expected, as the α5β1 integrin dimer is a major fibronectin binding receptor. IGF1R knockdown also failed to induce a significant increase in adhesion when cells were plated on fibronectin (Figure 6C-D), suggesting that fibronectin may promote adhesion regardless of IGF1R function.

**Figure 6.**
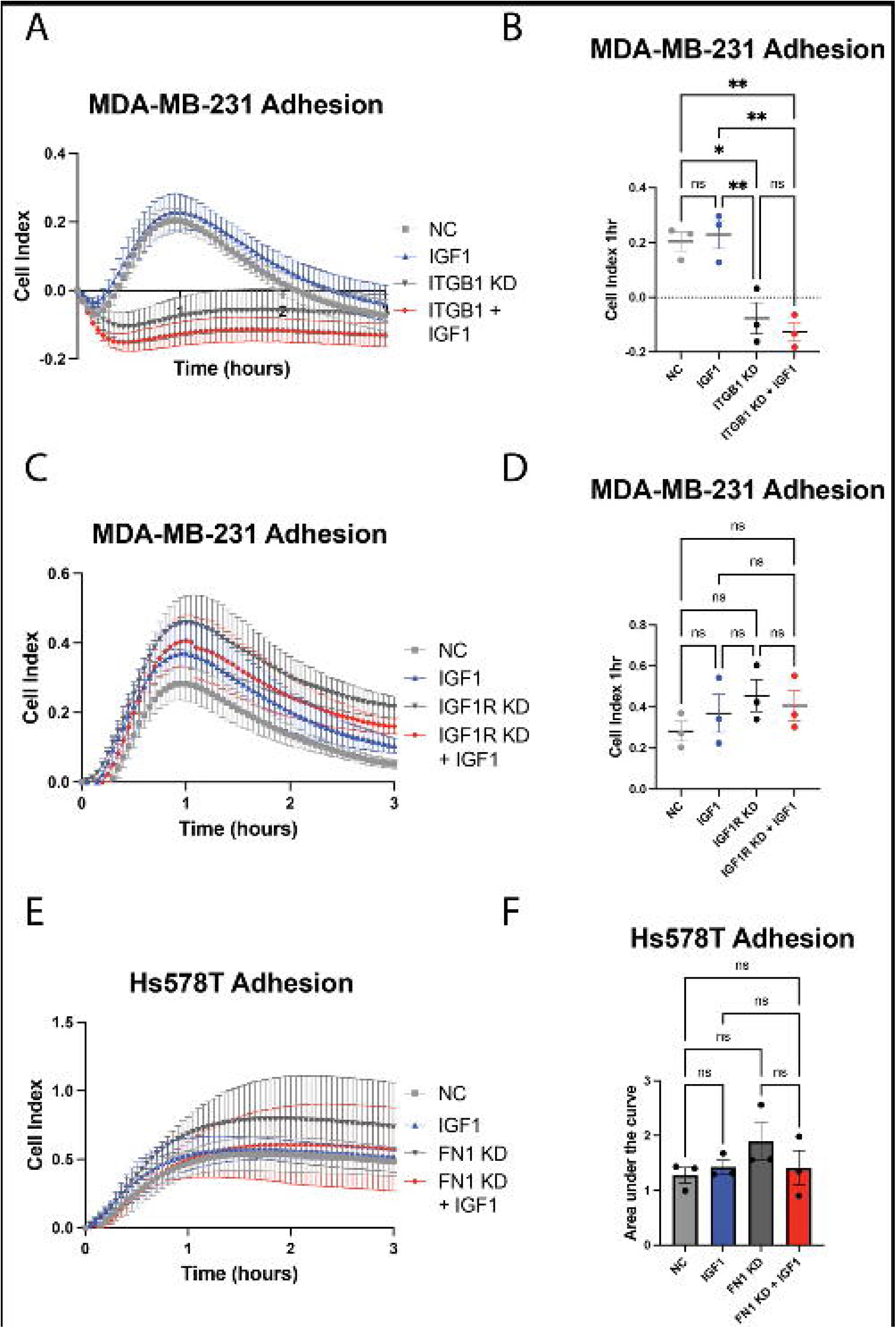
Stimulation with fibronectin inhibits IGF-1-mediated adhesion and promotes β1 integrin-dependent adhesion. A,C) Representative tracings from xCELLigence adhesion assay on E-plates coated with fibronectin prior to seeding MDA-MB-231 cells transfected with either anti-β1 integrin siRNA (A) or anti-IGF1R siRNA (C) versus a negative control (NC) siRNA. B,D) Quantification of adhesion from (A) and (C), respectively, from 3 independent experiments using peak adhesion values at 1-hour. E) Representative tracings from xCELLigence adhesion assay on uncoated E-plates to assess the effect of fibronectin siRNA knockdown on Hs578T cell adhesion, quantified in (F). *p<0.05, **p<0.005

Since the Hs578T cell line expresses fibronectin and is insensitive to IGF-1-mediated increases in adhesion, we were curious to determine if siRNA-mediated knockdown of fibronectin in these cells might sensitize them to IGF-1. FN1 knockdown was validated via western blot and qPCR (Supplemental Figure 5). As for our prior findings, IGF-1 stimulation had no effect on Hs578T adherence (Figure 6E, F). Fibronectin knockdown, whether in the presence or absence of IGF-1 stimulation, also did not alter Hs578T adhesion (Figure 6E, F), indicating that fibronectin expression by these tumor cells is not the reason they are insensitive to IGF-1-induced changes in adhesion.

To further understand the relationship between IGF-1 and fibronectin-induced cell adhesion we performed a second type of adhesion analysis by analyzing the number of cells attached to substrate on chamber slides after IGF-1 stimulation in the presence or absence of IGF1R inhibition with BMS (Figure 7A-D). On PDL–coated substrates, IGF-1 treatment significantly increased the number of attached cells compared to vehicle controls (Figure 7B), consistent with enhanced adhesion.

**Figure 7.**
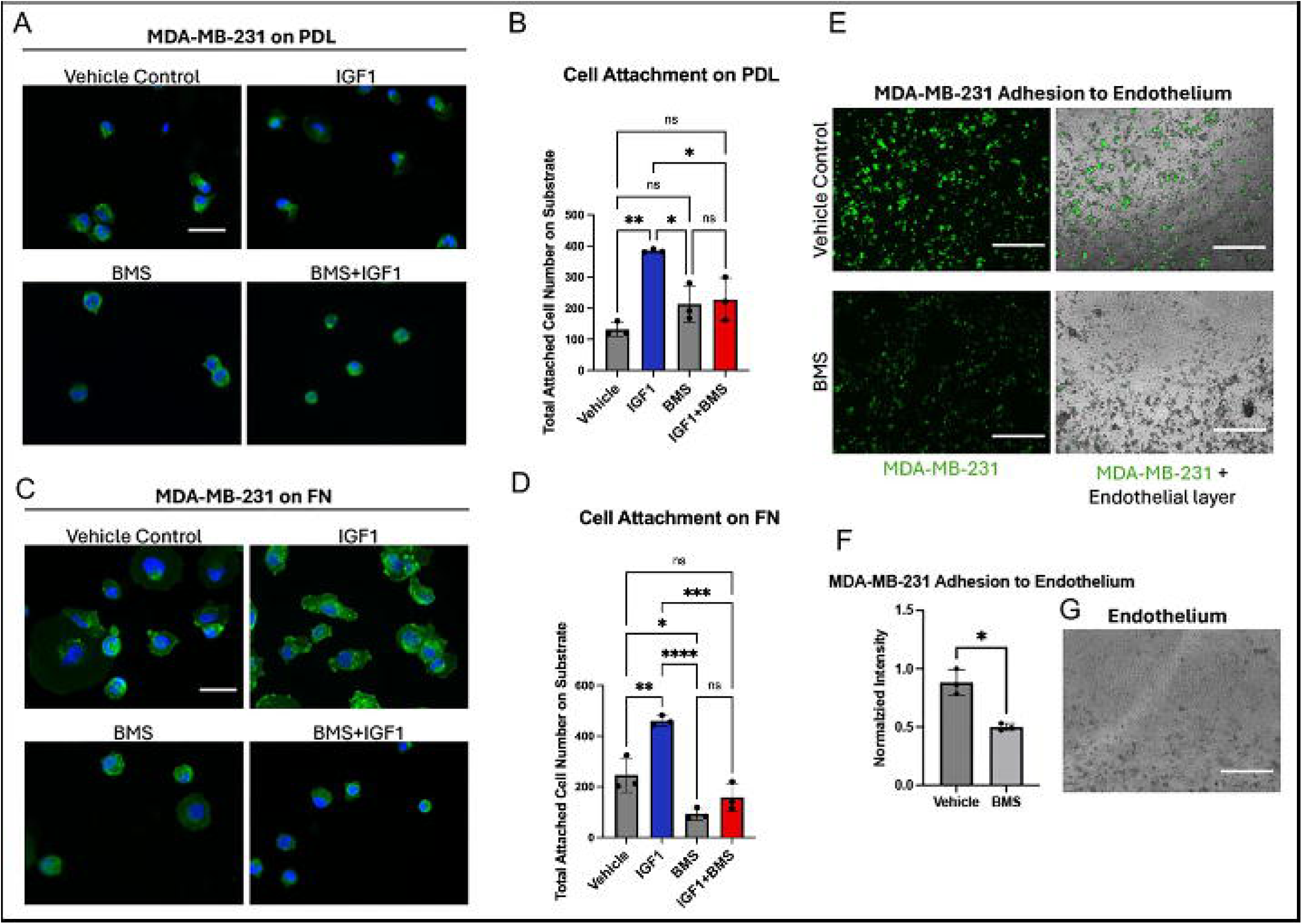
IGF-1 facilitates adherence to fibronectin and the endothelium, which is reversed by the IGF1R inhibitor BMS-752807. A, C) Slides were coated with A) PDL and C) fibronectin prior to seeding MDA-MB-231 cells. Cells were treated with BMS or left untreated, and with IGF1 or EtOH (vehicle control), followed by staining with anti-β1 integrin immunofluorescent antibody (green) and DAPI for nuclei (blue). Scale bar represents 50 µm. (B, D) Quantification of MDA-MB-231 cell attachment showing the total number of attached cells on substrates coated with B) PDL or D) fibronectin analyzed by one-way ANOVA and Tukey’s post hoc test. E) Green membrane dye– stained MDA-MB-231 cell attachment on the endothelial barrier after 24 h, with or without BMS treatment. Scale bar = 750 µm. F) Quantification of normalized fluorescence intensity showing attached MDA-MB-231 cells on the endothelial barrier after 24 h. G) Representative image of endothelial barrier formation after 24 h prior to MDA-MB-231 cell seeding. Scale bar = 750 µm. *p<0.05, **p<0.005, ***p<0.0005, ****p<0.0001.

Conversely, inhibition of IGF1R with BMS treatment markedly reduced cell attachment stimulated by IGF-1 (Figure 7B). Surprisingly, IGF-1 treatment similarly enhanced cell adhesion on fibronectin-coated substrates in this assay, which was inhibited by pre-treatment with BMS (Figure 7D).

Interestingly, BMS treatment also significantly reduced the number of cells adhered to fibronectin in the absence of IGF-1 stimulation (Figure 7D). The discrepancy between the findings in this experiment and the xCELLigence adherence assay with fibronectin may be due to way in which each assay quantifies cell adherence. xCELLigence measures adhesion using electrical impedance induced by cell coverage of the electrodes at the bottom of the E-plates wells. There is no wash-out step, so weakly adherent cells remain in the well and may contribute to electrical impedance and the resultant measure of adhesion. In the immunofluorescent assay, we specifically quantify of number of cells tightly adhering to substrate after a washout step and subsequent staining. As such, this assay is a more sensitive measure of strong cell attachment, indicating that IGF-1 promotes strong attachment of TNBC cells to fibronectin. Strong binding to fibronectin correlates with a more aggressive phenotype, underscoring the role of IGF-1 in promoting an invasive phenotype. Notably, the inhibitory effect of BMS appeared more pronounced on fibronectin than on PDL, suggesting substrate-dependent differences in integrin-mediated adhesion. Overall, these results demonstrate that IGF-1 enhances, while BMS suppresses, MDA-MB-231 cell attachment, and that fibronectin amplifies the sensitivity of cell adhesion to IGF1R inhibition.

Finally, to contextualize the effect of IGF-1 on adhesion in the broader cascade of metastasis we extended our analyses to tumor cell attachment to endothelial cells. Integrins are involved in multiple steps along the pathway to metastasis, and play a particularly crucial role in adhesion to the ECM as well as intravasation through the vascular endothelium (44). For instance, one study found that inhibition of melanoma cell binding to fibronectin reduced experimental metastases in C57BL/6 mice (45). To determine if IGF-1-dependent adhesion is involved in adhesion to vascular cells, we performed a binding assay to determine MDA-MB-231 cell attachment to HUVEC human endothelial cells. MDA-MB-231 cells were pretreated with a fluorescent dye for visualization and with BMS to inhibit IGF1R function prior to seeding on a layer of HUVEC cells. Treatment with BMS was terminated during the incubation period so that IGF1R expressed by HUVEC cells remained uninhibited. Strikingly, IGF1R inhibition significantly reduced MDA-MB-231 cell adhesion to HUVEC cells by 1.77-fold (Figure 7E-F). These results correlate well with the xCELLigence data from Figure 2, whereby BMS reduced MDA-MB-231 adhesion by approximately two-fold.

## Discussion

Our understanding of IGF1R function in cancer cell physiology is still evolving more than a decade after IGF1R inhibitor clinical trials began. In addition to its role in tumor cell proliferation and survival, there is mounting evidence that IGF1R also participates in tumor cell adhesion. Prior investigations have focused on the effect of integrin function on IGF1R signaling, but our data suggest that the relationship is reciprocal. We provide evidence that IGF1R regulates cell adhesion through β1 integrin and further, that this adhesion extends to binding the vascular endothelium.

The main function of integrins is to convey signals from the extracellular environment to the cell through attachment to the cytoskeleton. This interaction is bidirectional, and this two-way signaling axis has been dubbed “outside-in” and “inside-out” signaling (39). Several integrin inhibitors have been developed for various indications (see review by Pang and colleagues (39)). Integrins are an attractive therapeutic target against cancer in their own right, and expression of the β1 and β3 integrin subunits is associated with the transition of epithelial cancers to an aggressive, mesenchymal-like state (40). Integrin inhibitors have also been tested in cancer clinical trials; however, similar to IGF1R inhibitors, none have passed phase 3 (46).

Our findings thus far are summarized in a model presented in Figure 8, in which we propose that membrane-bound IGF1R inhibits β1 integrin function. We have shown that IGF-1 promotes adhesion in some, but not all, breast cancer cells, and this adhesion requires both IGF1R internalization and β1 integrin expression. Cell adhesion is an important process during multiple steps of the metastatic cascade, particularly migration, invasion and penetration of the vasculature. Despite the correlation of increased cell adhesion with AKT phosphorylation in MDA-MB-231 cells, inhibition of IGF1R internalization, which blocked IGF-1 induced cell adhesion, had no impact on AKT phosphorylation, suggesting an alternate mechanism that is also likely independent of ERK signaling.

**Figure 8.**
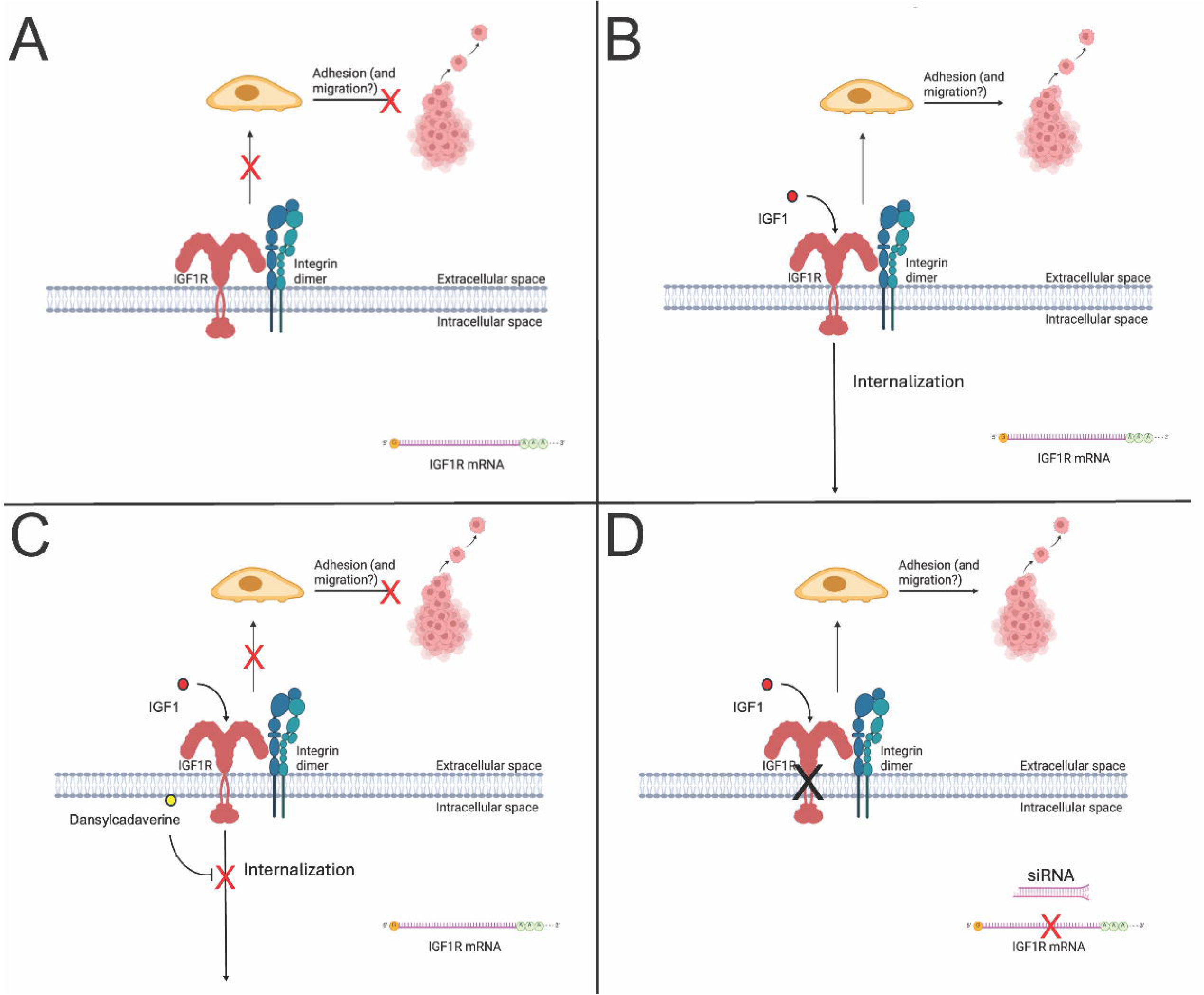
We hypothesize a model whereby membranous IGF1R protein inhibits β1 integrin function. A) In an unstimulated state, IGF1R is expressed on the cell surface and blocks adhesion mediated by β1 integrin. B) IGF-1 induces IGF1R internalization, which promotes integrin function and, as a result, adhesion. C) Inhibition of receptor internalization via dansylcadaverine inhibits IGF-1-mediated adhesion. D) IGF1R knockdown enhances cell adhesion mediated by β1 integrin. This figure was made with images from BioRender and edited with Microsoft PowerPoint. Galifi, C. (2025) https://BioRender.com/2c7ynqn.

The role of fibronectin in promoting cell adhesion through integrin activity is consistent with other studies. Green and colleagues demonstrated that either IGF-1 or fibronectin promoted blastocyst adhesion, as quantified by outgrowth area; however, both stimuli in tandem did not increase adhesion more than either alone, indicating that both IGF-1 and fibronectin increase cell attachment through an overlapping pathway. Since xCELLigence assays measure electrical impedance induced by cell spreading, our results via xCELLigence are consistent with this, as stimulating MDA-MB-231 cells with IGF-1 on fibronectin-coated E-plates did not promote adhesion above baseline. However, in our immunostaining experiment on fibronectin-coated chamber slides, IGF-1 increased the number of bound cells versus control after a washout step and subsequent staining. This suggests that IGF-1 promotes adhesion strength to fibronectin, although cell outgrowth may be unchanged.

These results suggest that IGF-1 stimulation, through IGF1R internalization, promotes β1 integrin-mediated cell adhesion of MDA-MB-231 TNBC cells. β1 integrin knockdown significantly reduced peak MDA-MB-231 adhesion to fibronectin over the course of 3 hours, indicating that β1 integrin is essential for strong binding to fibronectin. This is unsurprising and consistent with the literature, since α5β1 integrin is well understood to bind fibronectin and support adherence to the ECM (47). However, our visualization of cell binding to fibronectin on chamber slides suggests that IGF-1 treatment may further enhance cell binding to fibronectin. Whether this additional binding is through a common downstream effector is unknown.

It is also important to note that there are differences between the two TNBC cell lines we tested in their response to IGF-1. While MDA-MB-231 cell adhesion increased in response to IGF-1, Hs578T cell adhesion did not, despite both cell lines having similar levels of total β1 integrin protein. In addition, the proteins PP2A and RACK1, which are involved in IGF1R-integrin crosstalk (42), are not differentially expressed in the MDA-MB-231 cell line versus the Hs578T line. Parsing out the differences between cancers and their sensitivity to IGF-1 and their mechanisms for regulating adhesion is important, since IGF1R activation does not necessarily correlate to receptor expression levels (15). Our findings take this observation a step further by showing that IGF1R inhibition and IGF1R protein downregulation have entirely different effects on cell physiology. It is important to note that although the initial function of IGF1R inhibitors designed for clinical use is to reduce downstream signaling, IGF1R antagonists also downregulate receptor protein levels over time (7). In contrast, BMS-754807 and other tyrosine kinase inhibitors of IGF1R have not been shown to downregulate IGF1R protein at the cell surface (48). Indeed, ubiquitination of the β subunit of IGF1R and initiation of endocytosis requires tyrosine kinase activity. This is consistent with our proposed model whereby IGF-1 induces receptor internalization, resulting in increased adhesion through β1 integrin that is reversed by BMS-754807. In preclinical studies during which treatment courses are short, the effects of IGF1R inhibition may not reflect the long-term effect of receptor downregulation on cancer aggressiveness. This may in part explain the disparity between the results of mouse xenograft studies versus transgenic mouse models of IGF1R inhibition. IGF1R inhibitors have been shown to reduce tumor burden and improve survival in mouse xenograft models of various cancers (49); however, in transgenic mouse models of IGF1R attenuation or deletion, tumors were more aggressive (8–10, 50). Studying the long-term effects of IGF1R inhibition as well as IGF1R downregulation or deletion will be important to delineate compensatory effects that the cell undergoes, which may ultimately lead to more aggressive cancer phenotypes.

The role of cell adhesion in metastasis is complex and affected by several factors, including integrin expression, ECM composition, nutrient availability. Often, functional *in vitro* assays assess cell migration and invasion as a proxy for metastatic capability, but adhesion is a crucial component of migration, invasion, and intravasation that may be overlooked. Excitingly, we have found a link between β1 integrin-IGF1R crosstalk and a functional effect on cell binding to endothelial cells, suggesting that IGF1R function may regulate the process of TNBC intravasation and extravasation. Although the mechanism behind this process remains unclear, we hypothesize that IGF-1, by promoting β1 integrin function, can induce initiation of the intravasation process. Paradoxically, our data suggests that IGF-1 promotes TNBC adherence by removing IGF1R from the cell surface, as illustrated in Figure 8.

Our findings highlight the complexity of growth factor–integrin crosstalk in regulating tumor cell adhesion. Future studies may explore the interplay of IGF1R and integrins in 3D models and *in vivo* to determine if our findings correlate to intravasation and metastasis. In addition, in this study we focused particularly on fibronectin, but it is possible that other ECM ligands, such as collagen or laminin, are also involved in IGF-1-mediated adhesion. More work will be needed to explore the implications of our findings, but our results clearly implicate IGF1R in β1 integrin function in ECM and endothelial cell adhesion.

## Acknowledgements

We sincerely thank everyone involved in the development and completion of this project. We thank all other members of the Wood lab that helped with the daily activities of the lab, including Quan Shang, Marie L. Mather, and Divya Kantak. We thank Alison E. Obr for her expertise in various lab techniques and for laying the groundwork for the adhesion project. We thank Rosemary O’Connor, Niamh McDermott, and Stephen O’Shea for discussions and insights about our work. We thank members of the Cellular Imaging and Histology Core at Rutgers CINJ, including Luke Fritzky, for assistance with imaging and immunofluorescence. Finally, we thank Robert Wieder, Raymond B. Birge, Utz Herbig, Sergei V. Kotenko, and Pingping Hou for their expertise and insights on our project.

## Funding Statement

This project was funded by National Cancer Institute R01 CA204312 and New Jersey Commission on Cancer Research (NJCCR) Bridge Grant COCR23RBG003 to TLW, NJCCR Pre-doctoral Fellowship COCR23PRF013 to CAG and National Science Foundation Division of Chemical, Bioengineering, Environmental and Transport Systems 2426919 to AKM.

## Conflict of Interest

The authors declare no conflicts of interest.

## Author Contributions

CAG generated data and analyses and wrote the initial text draft of the manuscript. EDK, LFA, KM, KE, and SSS generated data and analyses; EDK and KM edited the manuscript. JJB made essential intellectual contributions regarding experimental design and technique throughout the project. AKM provided assistance with microscopy. TLW guided the entirety of this project from its conception and and design and edited the manuscript.

## Data Availability Statement

The datasets generated for this study will be deposited on OSF and the link will be made available upon acceptance and publication.

## Notes

### Competing Interest Statement

The authors have declared no competing interest.

