## Supplemental Figures for "Insulin-Like Growth Factor 1 Receptor Regulates Breast Cancer Cell Adhesion through Beta-1 Integrin"

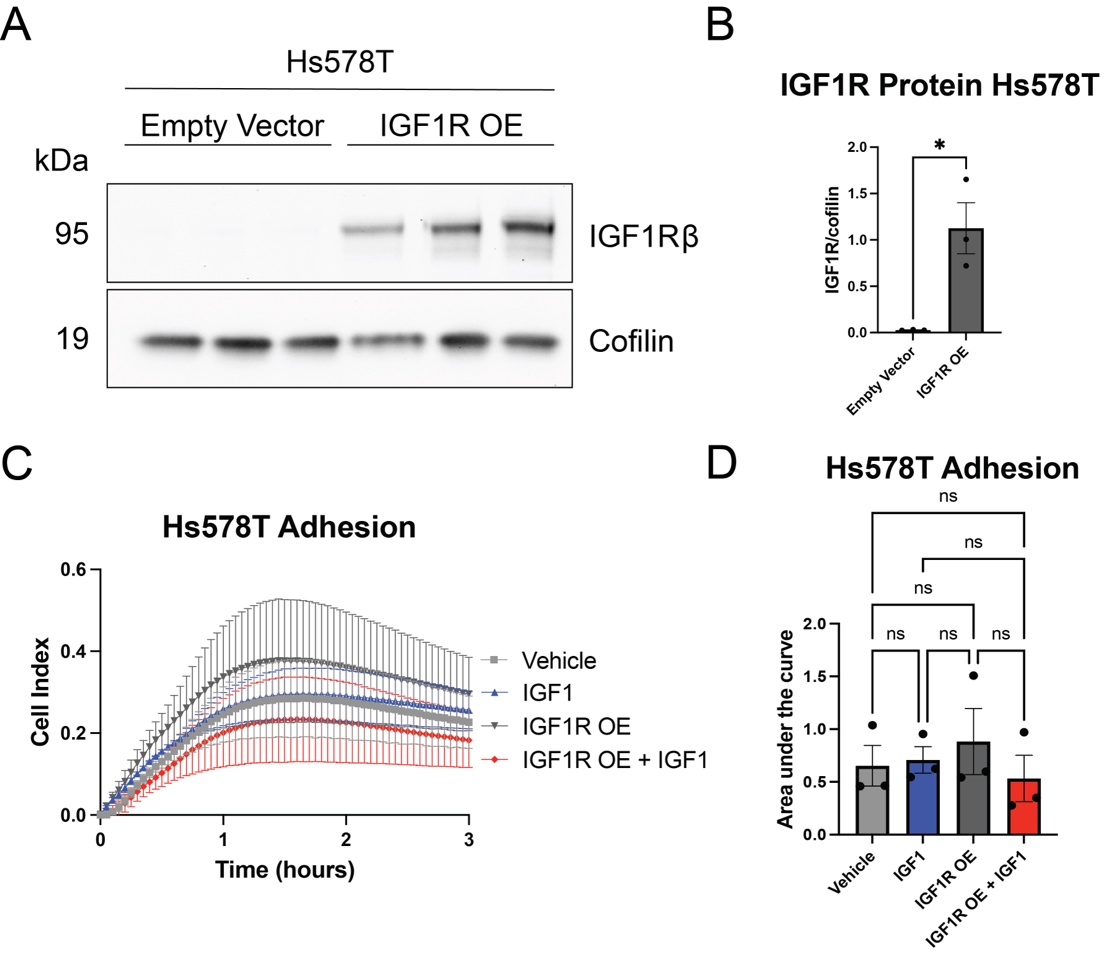

Supplemental Figure 1. IGF1R overexpression does not alter Hs578T adhesion in the presence or absence of IGF-1. A) Western blot validation of IGF1R overexpression after Lipofectamine 3000 plasmid transfection in Hs578T cells, B) Western blot data quantified by pixel intensity in ImageJ and analyzed for significance via one-tailed t-test. C) xCELLigence adhesion assay 2 days post-transfection. Cells were serum-starved for 3 hours and acutely stimulated with 10 nM IGF-1. D) Adhesion was quantified via area under the curve and groups were compared via one-way ANOVA and Tukey’s post hoc test. *p<0.05

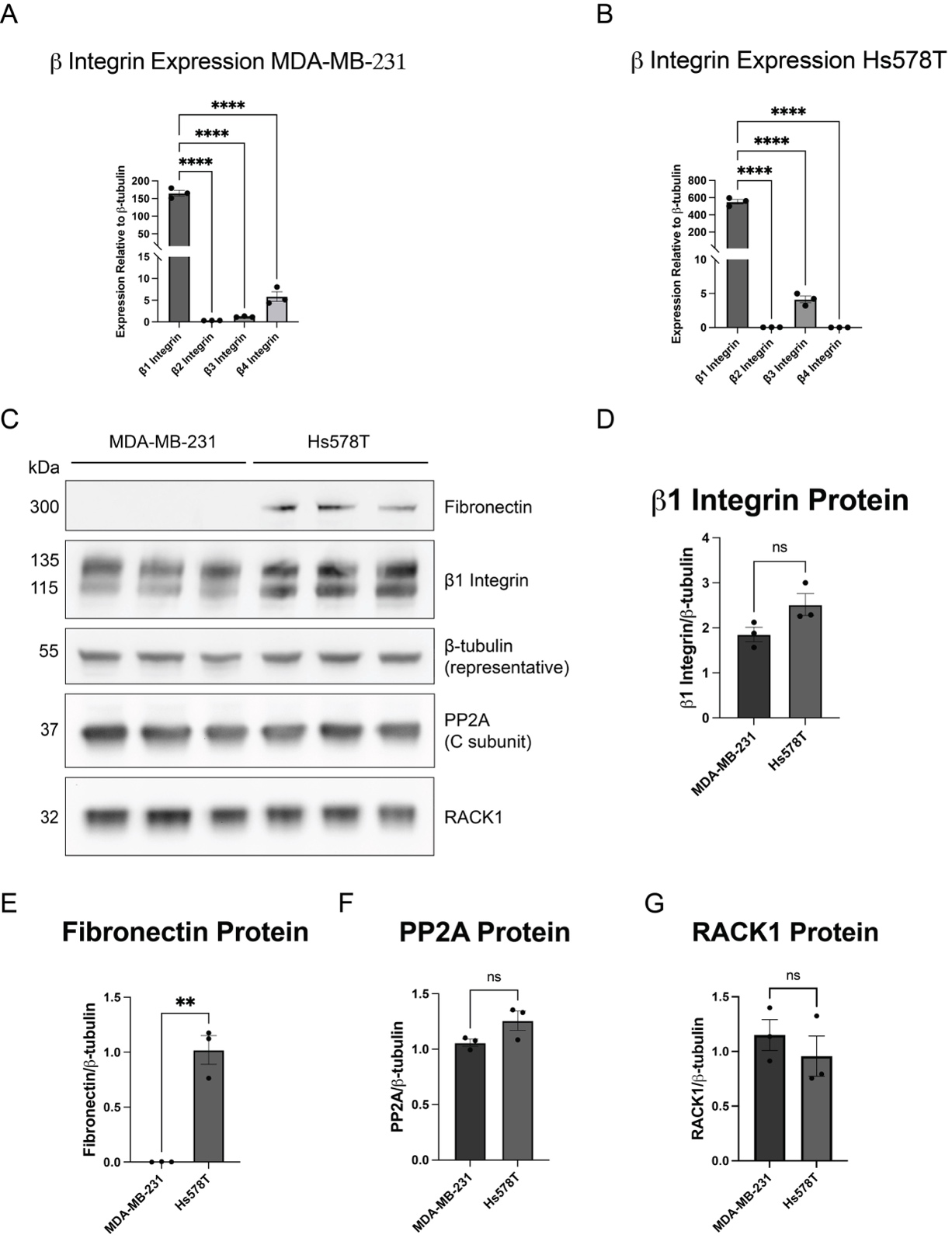

Supplemental Figure 2. β integrins and related proteins are expressed in TNBC cells. A) MDA-MB-231 and B) Hs578T cells were assessed for expression of integrins β1-4 via RT-qPCR. C) Protein expressions of fibronectin, β1 integrin, PP2A and RACK1 were analyzed in both cell lines via western blot and quantified in D-G). **p<0.005, ****p<0.0001

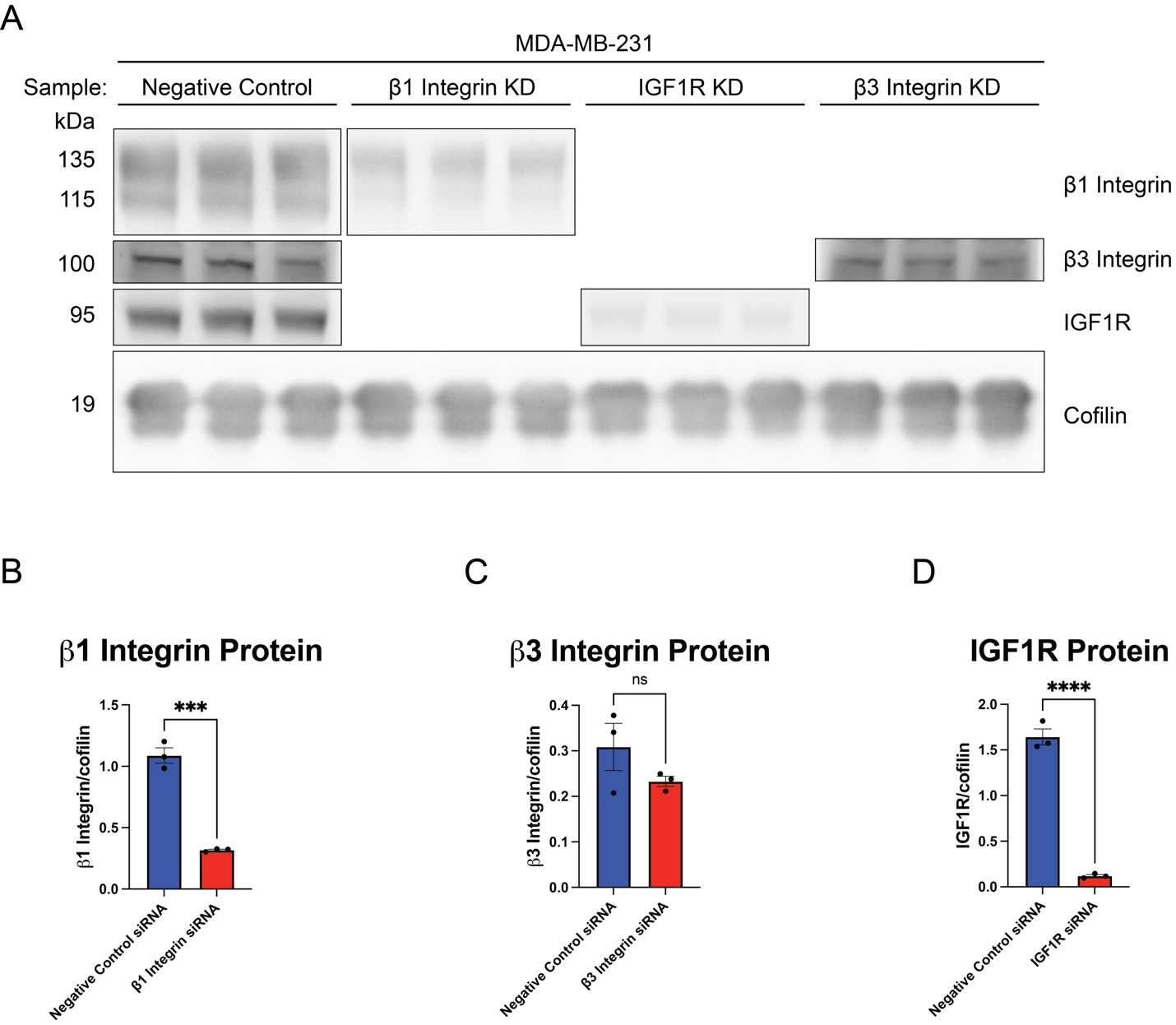

Supplemental Figure 3. Validation of anti-integrin and anti-IGF1R siRNA knockdowns in MDA-MB-231 cells. MDA-MB-231 cells were transfected concurrently with either a negative control siRNA (NC), anti-ITGB1 siRNA, anti-ITGB3 siRNA, or anti-IGF1R siRNA. A) Western blot showing corresponding bands for each protein. The 3 replicates for each group were separated by cutting the blot so that targets of similar size (ITGB1, ITGB3, and IGF1R) could be probed separately. Incubation and exposure times were kept consistent for each antibody in negative control and knockdown samples. B-D) Graphs showing quantification from Western blot. β1 integrin, β3 integrin, and IGF1R protein levels were compared between negative control and their respective knockdown groups and data were analyzed via two-tailed t-test. ***p<0.0005, ****p<0.0001

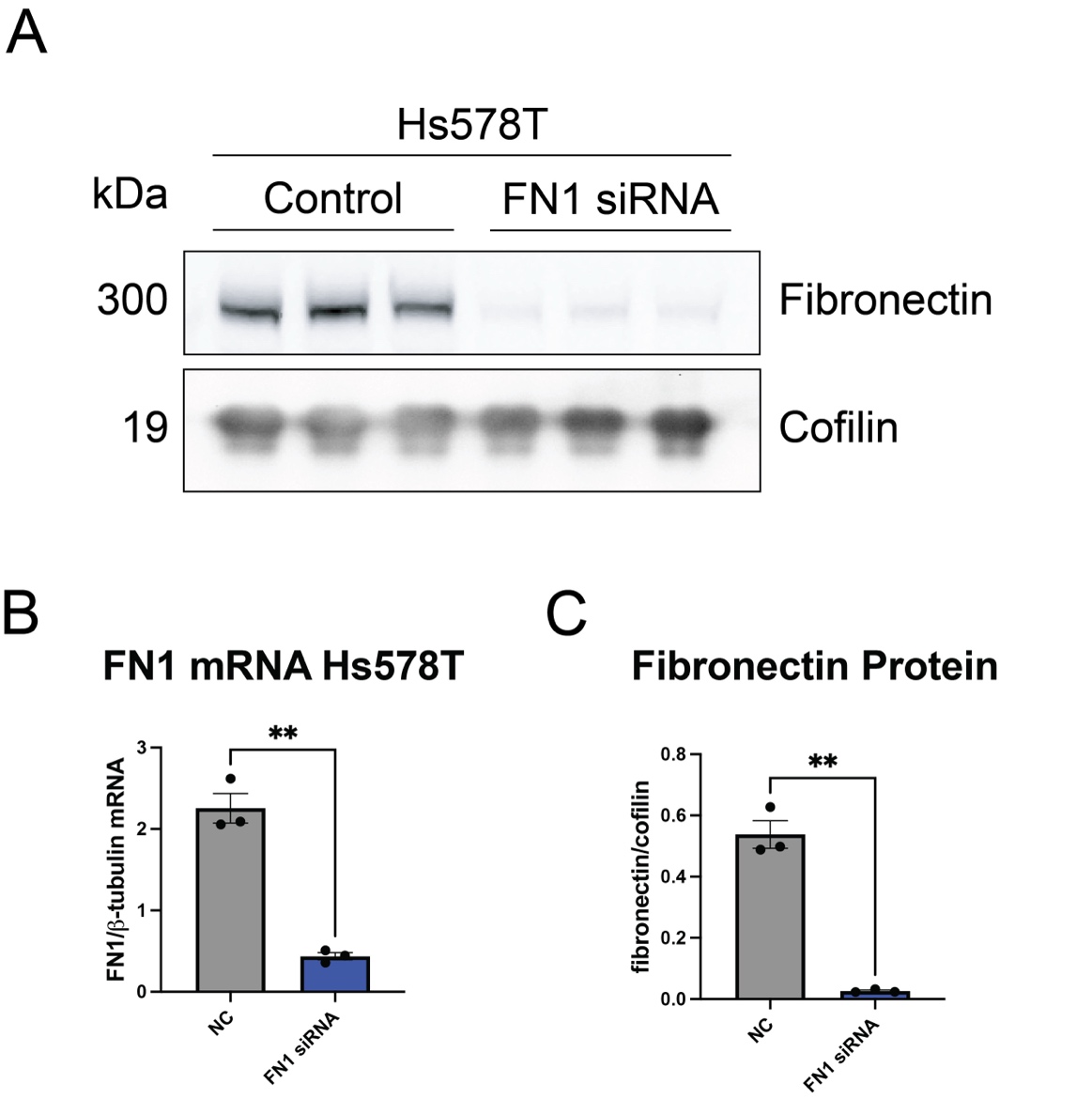

Supplemental Figure 4. Validation of FN1 knockdown in Hs578T cells via qPCR and western blot. Lipofectamine transfection was performed as outlined in the methods. A) Western blot showing fibronectin and the control, cofilin, quantified in C). B) qPCR was performed to determine FN1 mRNA knockdown versus β-tubulin mRNA.
